# Genome mining, phylogenetic, and functional analysis of arsenic (As) resistance operons in *Bacillus* strains, isolated from As-rich hot spring microbial mats

**DOI:** 10.1101/2022.03.20.485026

**Authors:** Aurora Flores, María F. Valencia-Marín, Salvador Chávez-Avila, Martha I. Ramírez-Díaz, Sergio de los Santos-Villalobos, Victor Meza-Carmen, Ma. del Carmen Orozco-Mosqueda, Gustavo Santoyo

## Abstract

The geothermal zone of Araró, México, is located within the trans-Mexican volcanic belt, an area with numerous arsenic (As)-rich hot springs. In this study, the draft genome sequence of two endemic *Bacillus* strains (ZAP17 and ZAP62) from Araró microbial mat hot springs was determined, which were able to grow on arsenate (up to 64 mM) and arsenite (up to 32 mM). Phylogenetic analysis based on 16S rRNA and *gyrB* sequences, as well as genome sequence analysis based on average nucleotide identity (>96%) and digital DNA–DNA hybridization (>70%), indicated that these strains belong to the *Bacillus paralicheniformis* ZAP17 and *Bacillus altitudinis* ZAP62. Furthermore, through genome mining, it was identified two arsenic resistance operons, *arsRBC,* and *arsRBCDA* in both strains as potential determinants of arsenic (As) resistance. Predicted ArsA (arsenial pump-driving ATPase), ArsB (Arsenical efflux pump protein), ArsC (Arsenate reductase), ArsD (Arsenical efflux pump protein) and ArsR (Metalloregulator/*ars* operon repressor) proteins, clearly grouped with their respective clades corresponding to other characterized bacterial species, mainly Firmicutes. To further evaluate the functionality of the *ars* operons in ZAP17 and ZAP62 strains, our results showed that *arsRBC* and *arsRBCDA* genes were expressed in the presence of arsenite (III). Finally, the presence of *ars* operons in the genome of *Bacillus* species residing in As-rich environments, such as the Araró hot springs, might be a potential mechanism to survive under such harsh conditions, as well as to design sustainable bioremediation strategies.

## 1. Introduction

Arsenic (As) is a widely distributed metalloid in both aquatic and terrestrial environments (Oremland and Stolz, 2003). Alternating among its various oxidation states, As is commonly found in nature bonds to oxygen, sulfur, and methyl groups, being arsenate (V) and arsenite (III) its most stable states (O’Day, 2006). Undoubtedly, geothermal fluids and volcanic emissions are important natural sources of arsenic since concentrations range from 0.01 to tens mg As / L ;around the world, this metalloid has been associated along boundaries of active plates (Wang et al., 2018). Is in this way, geothermal systems, such as hot springs. have been identified as natural sources of elevated As levels in water (Prieto-Barajas et al., 2017; Zhang et al., 2020).

In addition to being found naturally in the environment, human activities such as metal smelters, mining, and the use of pesticides have been associated as sources of As contamination in soil and groundwater, causing several human and animal health problems (Nriagu et al., 2007; Akopyan et al., 2018). Nowadays, the risk of poisoning and detrimental effects on human health from this metalloid has increased since elevated As concentrations have been found in groundwater (Zhang et al., 2019), exceeding the national guidelines of 0.01 mg / L for drinking (Ahmad and Bhattacharya, 2019).

As the most pathway for As movement into cells, arsenite, and arsenate manage to enter the cells through aquaglyceroporins (AQPs) and phosphate transporters (Pit) (Garbinski et al., 2019), once inside of cells causes harmful effects in living organisms (Mandal, 2017) since it induces oxidative stress by interfering in various cellular processes as it destabilizes proteins and causing may cause DNA damage to DNA (Lemire et al., 2013). However, bacterial and archaeal genomes throughout evolution have acquired genetic elements that allowed them to metabolize arsenic and participate in its cycling (Oremland and Stolz, 2003; Zhu et al., 2017). It is in this way that they achieved an autotrophy-based link with arsenic since the old anoxic atmosphere (Sforna et al., 2014).

There are exist, at least, four arsenic biotransformation mechanisms described in bacteria: (1) arsenic resistance system; (2) arsenic methylation and related pathways; (3) arsenite oxidation system; and (4) arsenate reduction system (Yan et al., 2019). This latter mechanism is widely distributed in prokaryotes, as a result of a selection process to survive in different environments, where the presence of this metalloid is widely abundant and distributed (Fekih et al., 2018). Arsenic resistance systems or *ars* systems are widely distributed in prokaryotes and these genetic elements are organized in *ars* operons, whose expressions are controlled by arsenite-responsive transcriptional repressor (ArsR) (reviewed in Fekih et al., 2018). It has been proposed that the transcriptional regulator ArsR was coded in conjunction with an arsenite efflux pump (ArsB) composing the simplest operon *arsRB* in the primitive atmosphere.Then, the change to an oxidizing environment must have led to the appearance of arsenate reductase enzymes (ArsC) and three-gene *arsRBC* operons since the majority of aqueous arsenite would have been oxidized to arsenate (Rosen, 1999; Mukhopadhyay and Rosen, 2002; Zhu et al., 2014; Fekih et al., 2018; Garbinski et al., 2019).

Due to the toxicity of this metalloid in the environment, it is important to investigate the genetic mechanisms behind the biotransformation processes of arsenic found in microorganisms, such as bacteria (Oremland and Stolz, 2003). Particularly, the Firmicutes are common inhabitants of aquatic systems, such as groundwater or hot spring microbial mats, where elevated concentrations of As constrain the survival of other bacterial species (Das et al., 2016; Prieto-Barajas et al., 2017; Zhang et al., 2020; Oremland and Stolz, 2003). Therefore, it is relevant to find out about the resistance mechanisms of this group of bacterial species that allows them to colonize and permanently be part of such microbial communities. Thus, the objectives of this work were (1) to sequence the genome of two *Bacillus* strains isolated from an As-rich hot spring and determine their arsenic resistance capacity; (2) to characterize the arsenic resistance systems through phylogenetic analysis and genome mining; and (3) evaluate their potential functionality under the presence of As (III).

## 2. Materials and Methods

### 2.1. Bacterial growth and Susceptibility tests

Strain ZAP17 was previously isolated and characterized from the geothermal system of Araró, México (Prieto-Barajas et al., 2017). Here, a new strain, ZAP62 was also isolated from a further screen to characterize highly As-tolerant strains from those microbial mats. These isolates were cultivated in Nutritive Agar (AN) culture medium, at 37 ° C, for routine use in the Laboratory and conserved at −70 ° C in 50% glycerol plus nutrient medium.

To determinate arsenite (As III) and arsenate (As V) resistance ability in ZAP strains, the bacteria were incubated in M9 medium at 37 ºC, 150 rpm for 12 h followed by diluting to a uniform optical density of 0.5 at 590 nm, before further inoculation. Aliquots of 500 µL of each bacterial suspension were added into M9 medium with 0, 4, 8, 12, 16, 20, 24, 28 and 32 mM of arsenite and 0, 8, 16, 24, 32, 40, 48, 56 and 64 mM of arsenate. After 24 h and 36 of incubation at 37 ºC and 150 rpm, the optical density was determined for ZAP17 and ZAP62 respectively.

### 2.2. DNA extraction and genome sequencing, assembly, and annotations

For DNA extraction, the strains were cultured on plates with nutrient agar from which a single colony was selected to be transferred to a liquid medium. The broths with the strains were incubated at 37 ºC overnight and were used for genomic DNA isolation. The genomic DNA was extracted using SDS/proteinase K and precipitating polysaccharides in the presence of high salt (Mahuku, 2004). In order to meet the quality standards required for sequencing, DNA samples were evaluated by agarose gel electrophoresis and by using a NanoDrop (Thermo Scientific).

The genomic DNA of ZAP strains was sequenced by Mr. DNA (Texas, USA,) using Illumina NovaSeq technologies. The assembly was performed using SPAdes assembler version v3.12.0 SPAdes (Bankevich et al., 2012). The draft genome of ZAP17 was built by using the reference chromosome replicon of *Bacillus paralicheniformis* ATCC 9945a (NCBI accession: CP005965.1), whilst the draft genome of ZAP62 was assembled using the genome of *Bacillus pumilus* TUAT1 (NCBI accession: AP014928.1). The final sequences obtained were deposited in the GenBank database and annotations were carried out by the NCBI Prokaryotic Genome Annotation Pipeline (PGAP) using the best-placed reference protein set method and GeneMarkS-2+. Final drafts genomes can be found with accession numbers: CP049698.1 and CP049589.1.

### 2.3 Taxonomic affiliation of the studied Bacilli strains

Phylogenetic analyses were performed based on 16S rRNA and *gyrB* gene sequences, which were extracted from ZAP17 and ZAP62 draft genome sequences obtained in this study, likewise, nucleotide sequences of related *Bacillus* strains were retrieved from NCBI GenBank. Sequences were aligned using the ClustalW algorithm, and trees were generated by Maximum likelihood (ML) using 1000 bootstrap replicates in MEGA 7.0.26 (Kumar et al., 2016).

Additionally, overall genome-related indexes (OGRIs) were used to identify the bacterial species of ZAP strains according to the proposed minimal standard (Chun et al., 2018). Briefly, the genome of ZAP17 and ZAP62 strains were evaluated for contamination usingContEst16S (https://www.ezbiocloud.net/tools/contest16s). Once contamination was not detected, the 16S rRNA sequence from these bacterial genomes were used for a taxonomic affiliation using the database of EzBioCloud (https://www.ezbiocloud.net/)(Yoon et al., 2017). Strains type that showed a ≥ 98.7% identity were chosen for species assignment using ANI (https://www.ezbiocloud.net/tools/ani) and dDDH (https://ggdc.dsmz.de/ggdc.php) algorithms.

### 2.4. Genome mining and phylogenetic analysis of *ars* operons

To detect the genetic elements involved in resistance to As, coding DNA sequences involved in this mechanism were manually searched through blast and genomic annotations. Likewise, to have an overview of the evolutionary status of *ars* genes phylogenetic trees were constructed by using protein sequences that were blasted against Non-redundant RefSeq proteins and UniprotKB/Swiss–Prot in the NCBI database. Phylogenetic trees were built by using the ML method in MEGA 7.0. 26 with 1000 bootstrap replicates (Kumar et al., 2016).

### 2.5. Gene expression analysis

To determine if arsenic resistance elements were functional, a genes expression assay was performed. For this induction assay, the strains were incubated in M9 supplemented with 0 or 12 mM. Each treatment was established in duplicate for each strain. After 24 h of incubation at 37 ºC and 150 rpm, the optical density was determined, and the total RNA was extracted using TRIzol reagent from Invitrogen according to the manufacturer’s instructions. After extraction, the RNA underwent a digestion process to remove DNA from the samples before PCR applications. This was accomplished using RQ1 RNase-Free DNase purchased from Promega and following the supplier’s instructions. After the enzymatic digestion process, the RNA samples were used for cDNA synthesis, this was achieved using the GoScript Reverse Transcription System kit from Promega, following the manufacturer’s instructions. The PCR reactions were achieved by using cDNA obtained from strains growing at 0 or 12 mM of arsenite, and a different set of primers, which were designed to amplify the arsenic resistance genes of the ZAP strains and 16S rRNA genes (Table 1). The mentioned primers were designed using the genomes of strains in Primer-BLAST (https://www.ncbi.nlm.nih.gov/tools/primer-blast/), and tested using genomic DNA of each strain as a template as well as the cDNA obtained from each treatment in conjunction with Promega’s GoTaq Green Master Mix.

**Table 1.**
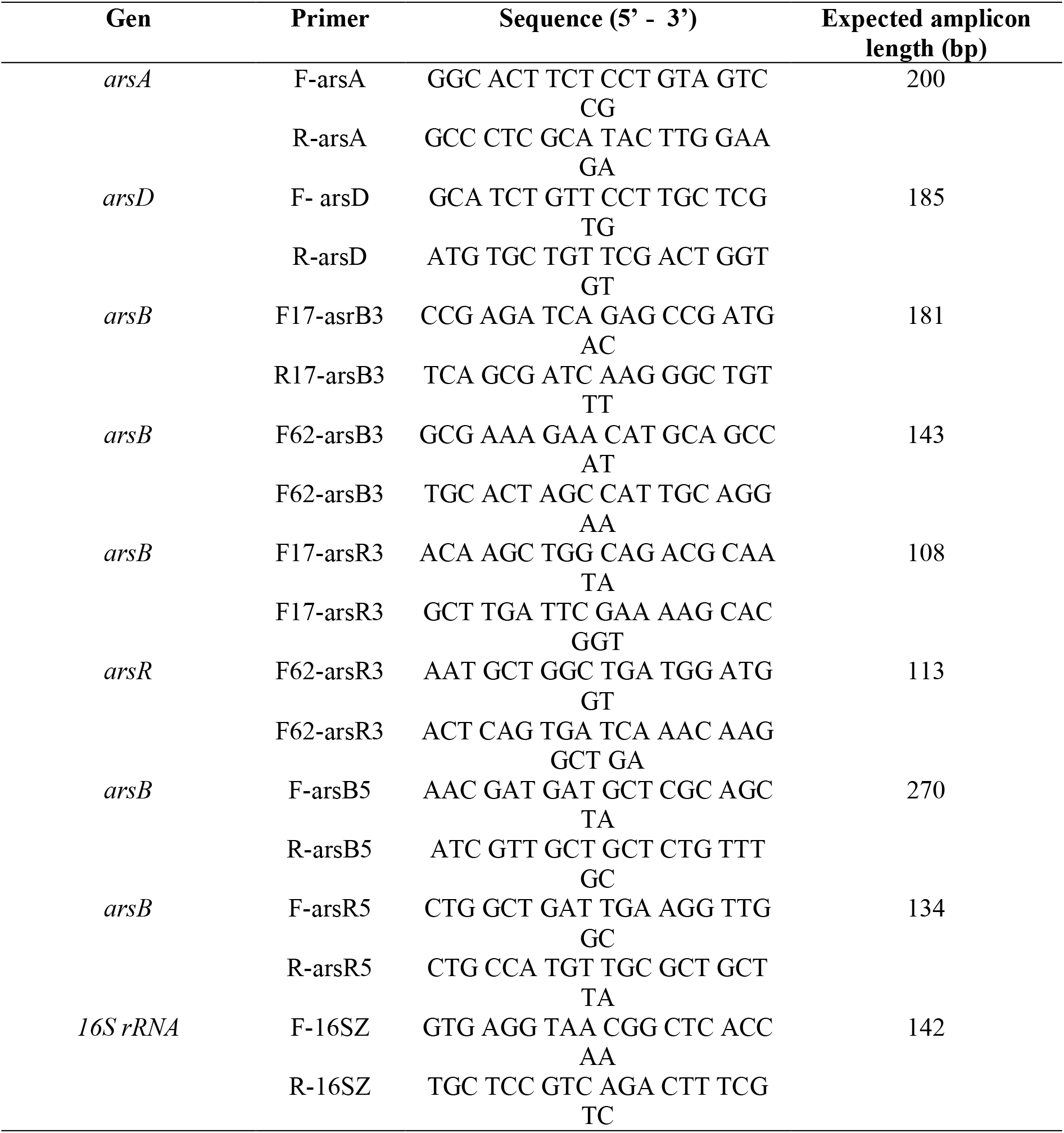
Primers designed and utilized in this study.

### 2.6. Statistical analysis

Data are presented as means with standard error. Significant differences were determined according to Duncan’s test using an alpha of 0.05.

## 3. Results and discussion

### 3.1. Evaluation of As tolerance

In this work, As-tolerance was evaluated in two bacterial strains, ZAP17 and ZAP62, which were isolated from As-rich hot spring microbial mats environments (Prieto-Barajas et al., 2017). Figure 1 shows arsenite (As III) tolerance in upper panels of strains ZAP17 (a), and ZAP62 (b); bottom panels represent arsenate (As V) tolerance of ZAP17 (c), and ZAP62 (d). As expected, arsenate tolerance is bigger compared to growth in media supplemented with arsenite. In fact, both strains manage to resist up to 124 mM arsenate (with data showing only up to 64 mM), while their growth in medium with arsenite showed a drop in growth at 16 and 24 mM in ZAP17 and ZAP62, respectively. Other works have reported resistance to arsenate 5-10 times higher than concentrations of arsenite (Dunivin et al., 2018; Han et al., 2019). According to Botes et al. (2007), arsenic hyper-resistant bacteria can grow in 15 mM arsenite and up to 500 mM arsenate. Additionally Badage et al. (2020) undoubtedly showed that hypertolerance goes further since the isolate identified as *Bacillus firmus* L-148 can grow up to 3.3 M of arsenite and 4 M of arsenate.

**Figure 1.**
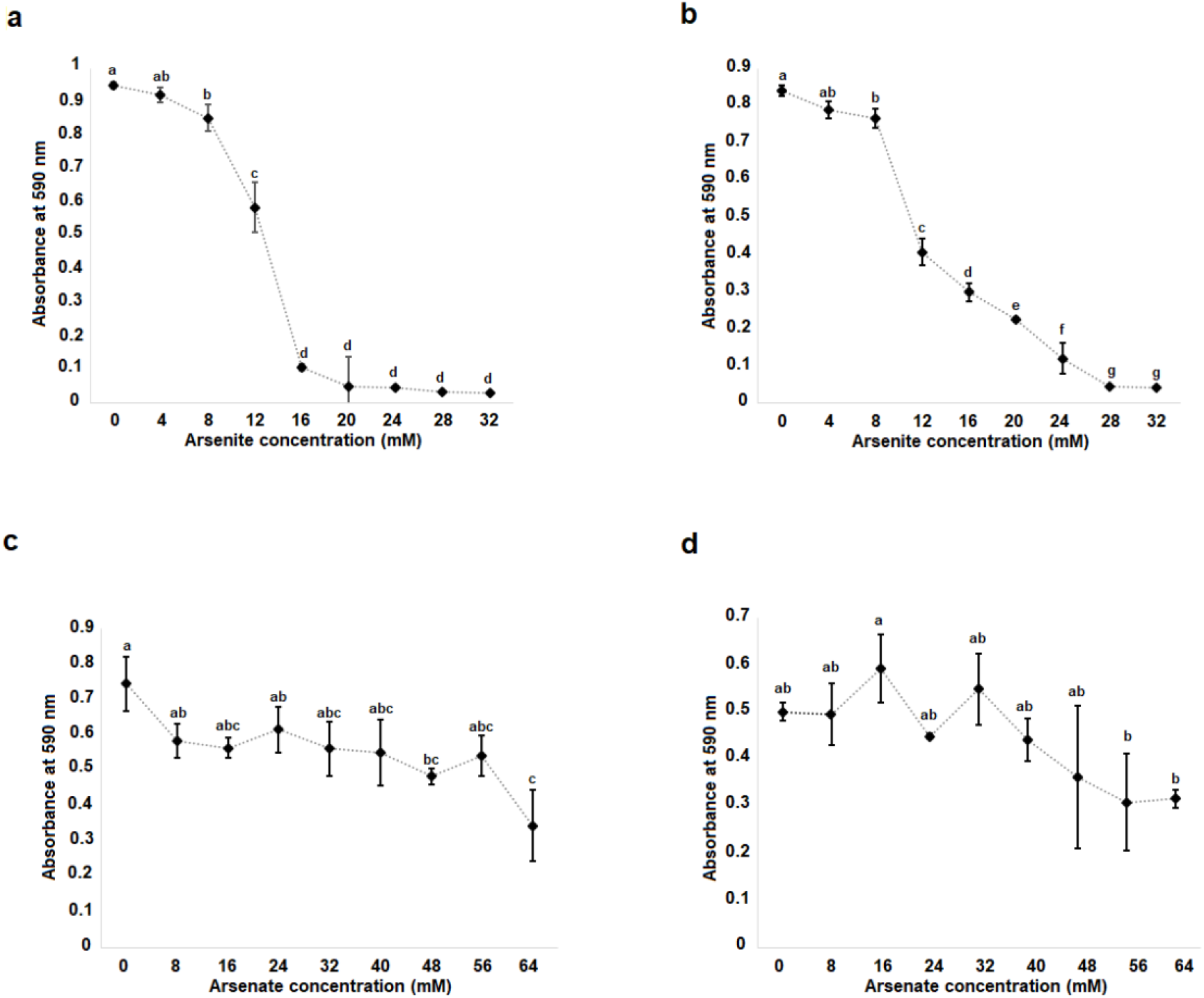
Arsenic resistance evaluation. Top panels represent arsenite resistance in ZAP17 (a), and ZAP62 (b); bottom panels represent arsenate resistance in ZAP17 (c), and ZAP62 (d). Data are presented as mean with standard error, significant differences were determined according a Duncan’s test using an alpha of 0.05, and are represented with different letters.

In a previous study from our group, the tolerance to arsenic was evaluated in a poor nutrient (M9) and rich (NA) media, observing elevated tolerance to arsenite and arsenate metalloids in NA compared to M9 (Prieto-Barajas et al., 2018). This is because NA contains higher amounts of phosphates (PO_4_^3-^), which can compete with arsenate, to be internalized through phosphate transporters (Rosen and Liu, 2009). Therefore, here we decided to analyze the tolerance of the strains only in a liquid and poor medium to not underestimate the As tolerance. Even so, tolerance to both forms of arsenic in strains ZAP17 and ZAP62 was higher than other As-tolerant strains of the *Bacillus* genus (Jia et al., 2019). Additionally, these levels of tolerance to arsenic are sufficient to survive in environments rich in As, such as the hot springs of Araró (Prieto-Barajas et al., 2017).

### 3.2. Genome sequencing and species affiliation

To have the genomic background of strains ZAP17 and ZAP62 associated with their tolerance to arsenic, their genomic DNA was sequenced and their closest taxonomic affiliation to bacterial species was carried out. Figure 2 (panel a and b) shows the micrographs of the strains ZAP17 and ZAP62, observing the bacilli form in both cultures. The chromosome size was 4.5 Mb, a % GC content of 45.80, and 4,690 genes for strain ZAP17. In the case of strain ZAP62, its chromosome had a size of 4.04 Mb, a % GC of 40.90, and 4,275 putative genes (Figure 2, paper c and d, and Table 2).

**Figure 2.**
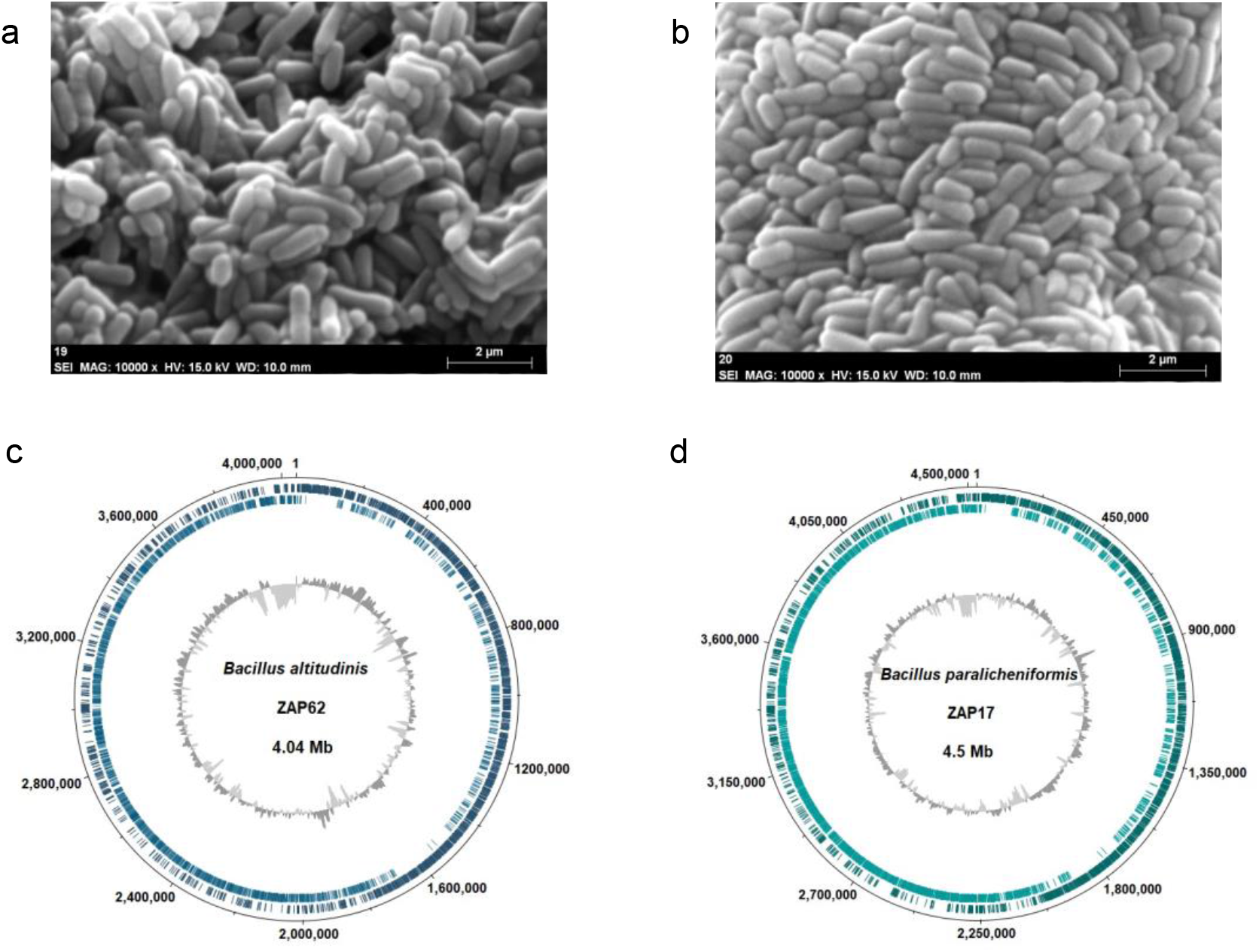
Scanning electron microscope photography of ZAP17 (a) and ZAP62 (b), and chromosome genome maps of ZAP17 (c) and ZAP62 (d). Dark green and light green tracks indicate genes on forward and reverse strand, respectively. The within-genome GC content variation is indicated by gray tones.

**Table 1.** Overview of genomic features of strains ZAP17 and ZAP62.

| **Attributes** | **ZAP17** | **ZAP62** |
| --- | --- | --- |
| Genome size (bp) | 4,528,645 | 4,041,532 |
| % GC | 45.80 | 40.90 |
| Number of contigs | 89 | 30 |
| N50 (bp) | 104,986 | 421,415 |
| L50 | 12 | 5 |
| Sequencing technology | Illumina NovaSeq | Illumina NovaSeq |
| Annotation pipeline | NCBI Prokaryotic<br>Genome Annotation<br>Pipeline (PGAP) | NCBI Prokaryotic<br>Genome Annotation<br>Pipeline (PGAP) |
| Genes (total) | 4,690 | 4,275 |
| CDSs (total) | 4,609 | 4,198 |
| Genes (coding) | 4,475 | 4,156 |
| Genes (RNA) | 81 | 77 |
| rRNAs | 1, 1, 1 (5S, 16S, 23S) | 1, 1, 1 (5S, 16S, 23S) |
| Complete rRNAs | 1, 1, 1 (5S, 16S, 23S) | 1, 1, 1 (5S, 16S, 23S) |
| tRNAs | 73 | 69 |
| ncRNAs | 5 | 5 |
| Pseudogenes | 134 | 42 |
| Genebank accession number | CP049698.1 | CP049589.1 |

The taxonomic affiliation of strains ZAP17 and ZAP62 was explored by using the 16S rRNA gene (Joung and Côté, 2002; Kim and Chun, 2014; Woese, 1987), and the *gyrB* gene, which is important to discriminate species in members of Bacilli (La Duc et al., 2004; Wang et al., 2007). Thus, these genes were able to distinguish the taxonomic and phylogenetic relationship of the studied strains, since there is a clear distinction ofZAP17 and ZAP62 and related to the groups of *Bacillus subtilis* and *Bacillus pumilus*, respectively (Figures 3 and Figure 4). However, the topologies of these phylograms are not entirely conclusive since only the tree using the *gyrB* gene showed a connection between ZAP17 and the species *Bacillus paralicheniformis* with a percentage greater than 85% of bootstraps.

**Figure 3.**
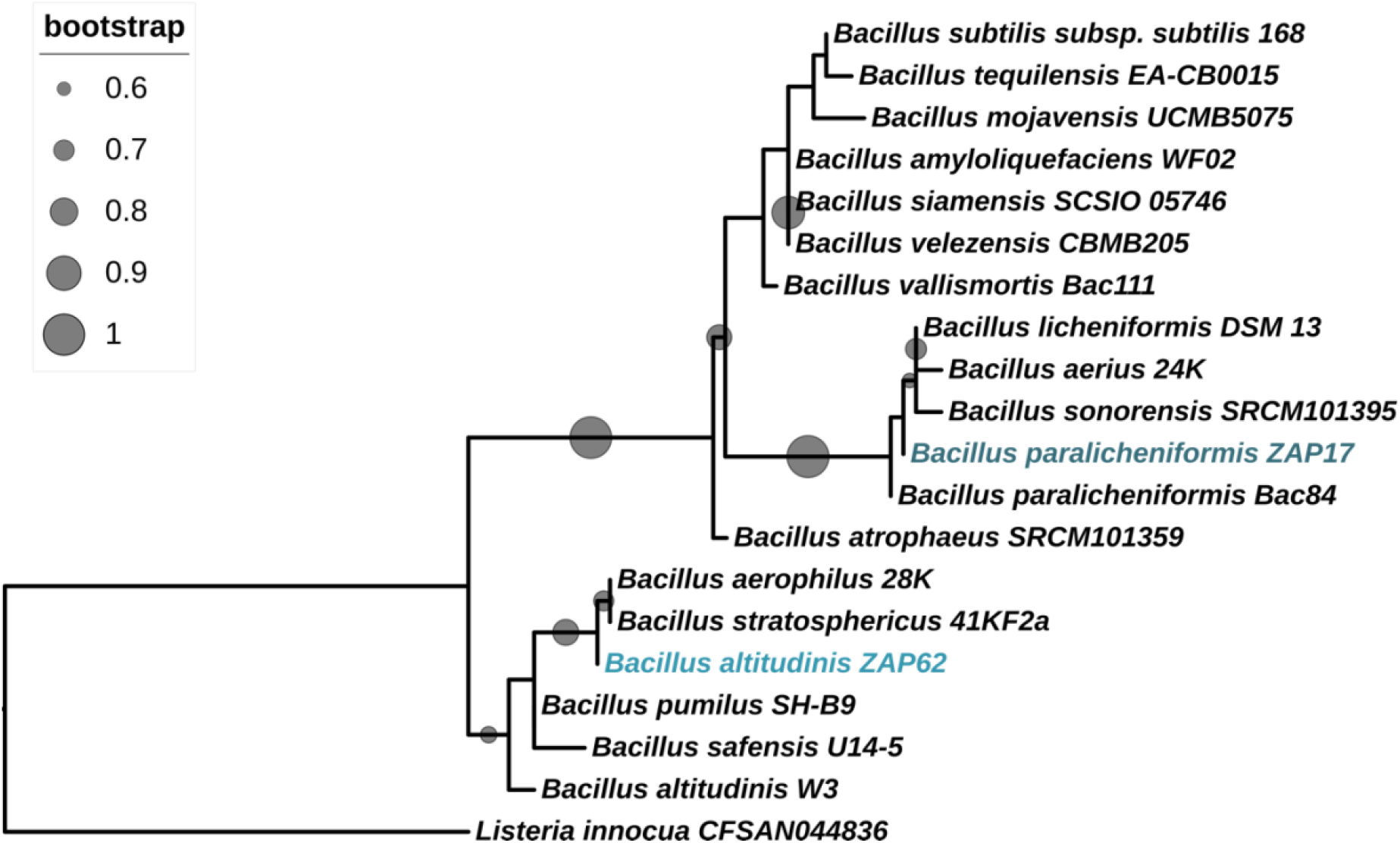
Phylogenetic relationships of ZAP strains with *Bacillus subtilis* and *Bacillus pumilus* groups inferred by analysis of *16S rRNA.* The phylogenetic tree was constructed using the Maximum Likelihood method and bootstrap support of 1000 replications. The inner circles show replications greater than 60%.

**Figure 4.**
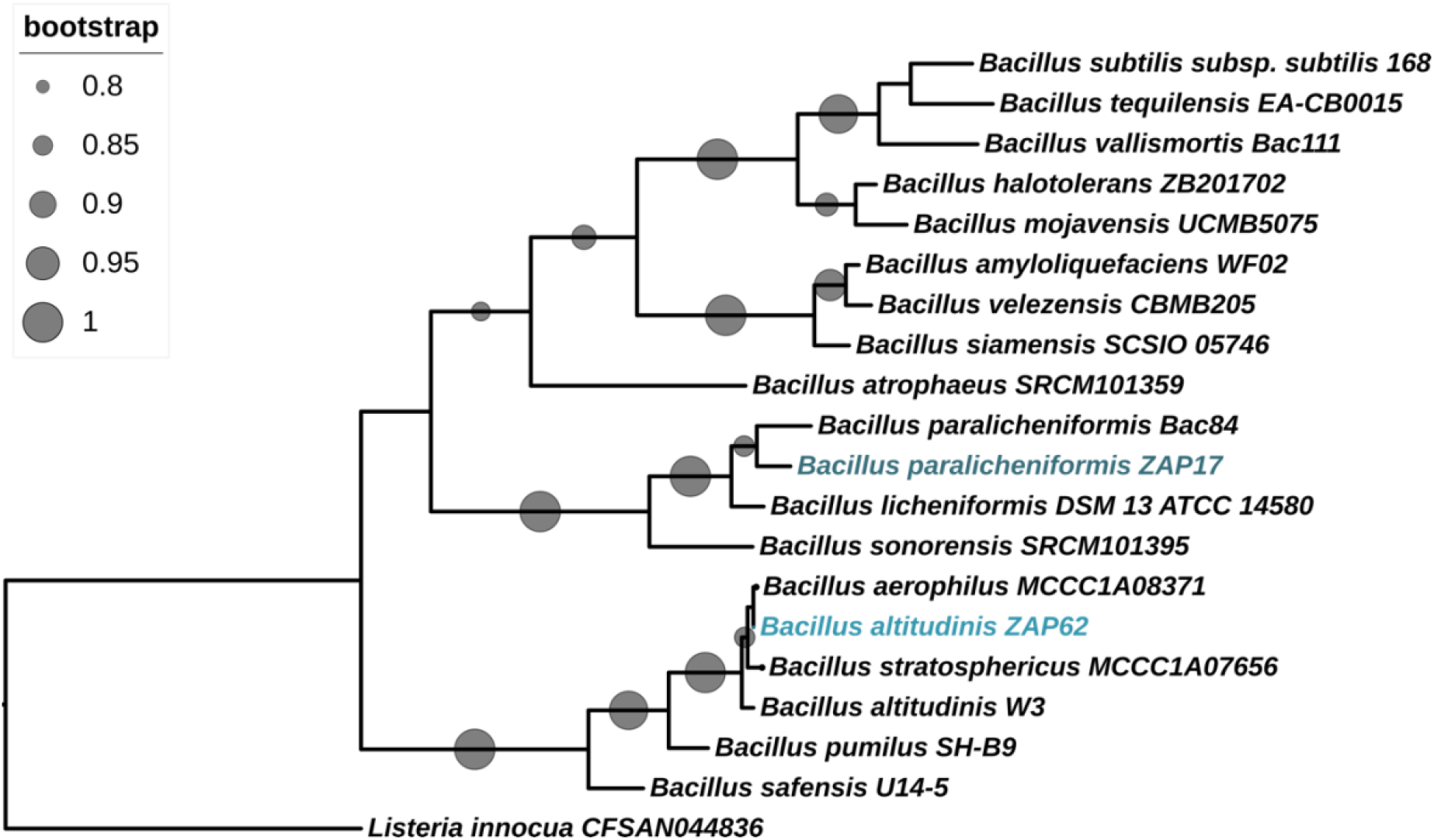
Phylogenetic relationships of ZAP strains with *Bacillus subtilis* and *Bacillus pumilus* groups inferred by analysis of *gyrB*. The phylogenetic tree was constructed using the Maximum Likelihood method and bootstrap support of 1000 replications. The inner circles show replications greater than 80%.

Therefore, it was additionally employed the overall genome-related indexes (OGRIs) methodology to solve the taxonomic affiliation of the ZAP strains, proposed by Chun et al. (2018). Thus, OGRIs revealed that strain ZAP17 belongs to the *B. paralicheniformis* since both ANI and DDH exceed the cutoff values (≥96% and ≥ 70% identity, respectively) for species determination (Table 3). Likewise, the average nucleotide identity and the percentage of *in silico* hybridization showed that the ZAP62 strain was highly affiliated to the *B. altitudinis* (Table 4).

**Table 3.** OGRIs values obtained from the comparison of ZAP62 and closely related species.

| **Species/Strain** | **16S**<br>**≥ 97 %** | **ANI**<br>**≥ 96 %** | **GGDC**<br>**≥ 70 %** |
| --- | --- | --- | --- |
| *Bacillus altitudinis* W3<sup>T</sup> | 99.93 | 98.47 | 86.6 |
| *Bacillus xiamenensis* VV3<sup>T</sup> | 99.86 | 91.43 | 43.9 |
| *Bacillus pumilus* SH-B9<sup>T</sup> | 99.59 | 88.76 | 36.1 |
| *Bacillus safensis* FO-36b<sup>T</sup> | 99.52 | 89.04 | 36.4 |
| *Bacillus atrophaeus* SRCM101359<sup>T</sup> | 97.48 | 70.85 | 17.9 |
| *Bacillus siamensis* SCSIO 05746 | 97.28 | 70.75 | 18.5 |
| *Bacillus velezensis* CBMB205<sup>T</sup> | 97.22 | 70.81 | 18.2 |
| *Bacillus subtilis* NCIB 3610<sup>T</sup> | 97.21 | 70.98 | 18.1 |
| *Bacillus amyloliquefaciens* DSM7<sup>T</sup> | 97.01 | 70.48 | 18.1 |
| *Bacillus tequilensis* EA-CB0015<sup>T</sup> | 97.01 | 70.99 | 18.1 |
| *Bacillus halotolerans* ZB201702<sup>T</sup> | 97.01 | 71.09 | 18.4 |

*B. altitudinis* has been isolated from various environments, including thermal pools (Abdollahi et al., 2021). Likewise, this species has been associated with good capacities to tolerate heavy metals, as well as excellent capacities to bioremediate environments contaminated with zinc, nickel, and copper (Khan et al., 2021; Babar et al., 2021; Yue et al., 2021). On the other hand, *B. paralicheniformis* KNPhave shown a good capacity to reduce chromate through three possible chromate efflux transporters. When analyzing the genome of strain KNP, genes that code for proteins involved in the reduction and resistance to As were also detected, such as thioredoxin, *arsA,* and *arsB* (Arora et al., 2021). Other strains of *B. paralicheniformis* have also been isolated from Espinazo Hot springs in Nuevo Leon, México (Silva-Salinas et al., 2021), Morocco (Maski et al., 2021), and Algeria (Benammar et al., 2020). Interestingly, all these *B. altitudinis* strains were associated with diverse biotechnological applications. However, to our knowledge, this would be the first work where the capacity of *B. paralicheniformis* and *B. altitudinis* species with good tolerance to metalloids such as arsenic is evaluated. Current investigations are being carried out to know its potential in the remediation of environments contaminated with such metalloids.

### 3.3. Analysis of the *ars* operons in ZAP17 and ZAP62 genomes

To better understand the associated mechanisms of As resistance, *ars* genes were identified in ZAP17 and ZAP62 genomes. Among all coding DNA sequences (CDS), two putative *ars* operons were found in both genomes and compared the synteny with other bacterial genomes, including several Bacilli species (Figure 5). One operon in ZAP strains consisted of 3 genes (*arsRBC*) and the second of 5-gene cluster *arsRBCDA*. The products of operon *arsRBC* in ZAP17 with locus tags G3M81_RS10920 (115 aa), G3M81_RS10915 (431 aa), and G3M81_RS10910 (139 aa) showed 100 % of amino acids identity compared with ArsR, a winged helix-turn-helix transcriptional regulator, ArsB, an arsenical efflux pump membrane protein, and ArsC, an arsenate reductase, proteins of *B. paralicheniformis.* Likewise, the products of *arsRBCDA* with locus tags in ZAP17 G3M81_RS14695 (139 aa), G3M81_RS14690 (432 aa), G3M81_RS14685 (139 aa), G3M81_RS14680 (125 aa), and G3M81_RS14675 (589 aa) showed high percents of identity not only with members of *B. subtilis* group such as *B. paralicheniformis* and *B. licheniformis*, but also with species as *B. altitudinis*, *B. swezeyi* and *B. dafuensis*.

**Figure 5.**
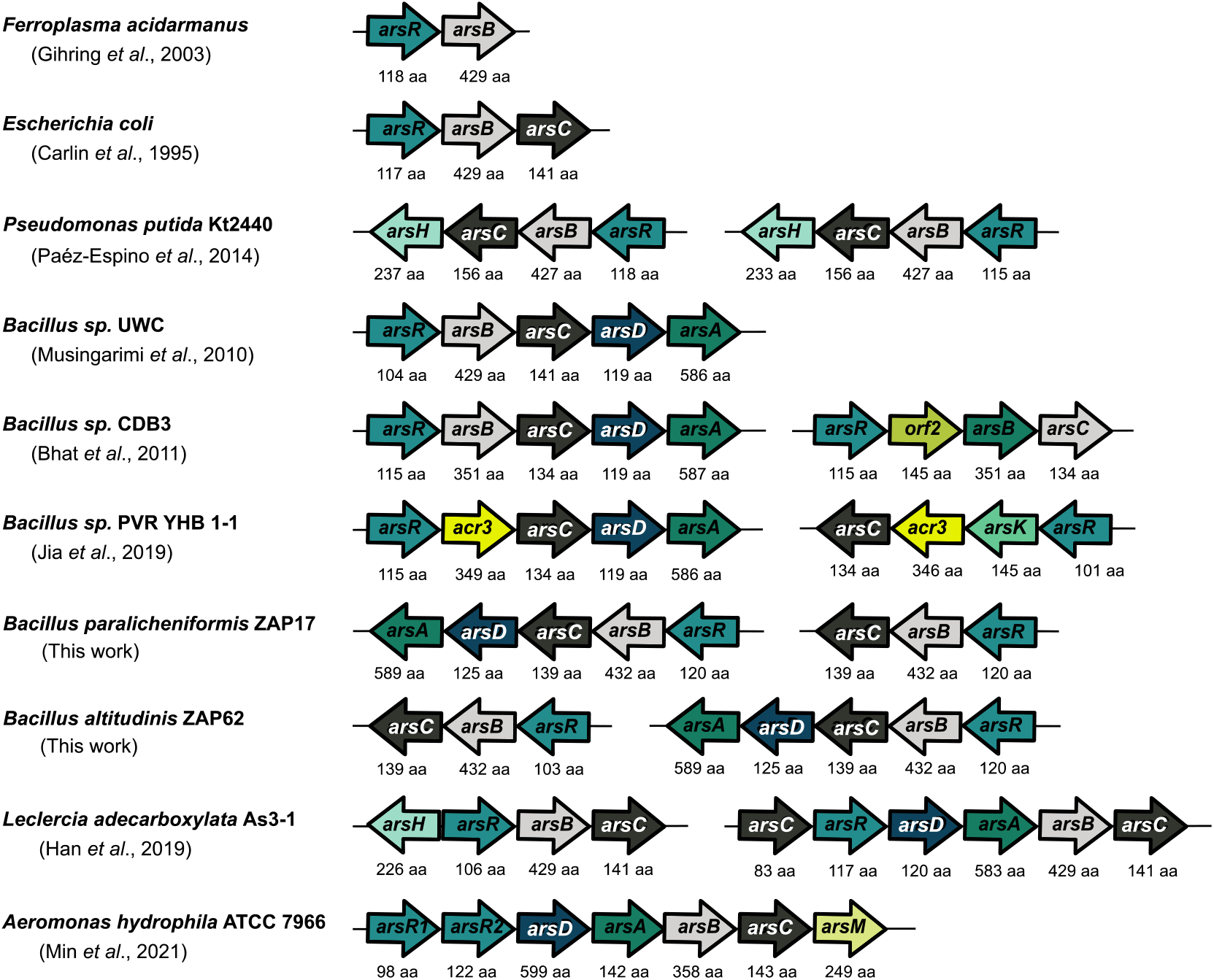
As resistance operons in ZAP strains and their comparison with other arsenic-resistant prokaryotes. Genetic organization of *ars* operons of some arsenic-resistant prokaryotes and their comparison with ZAP strains. Arrows with different colors represent open reading frames and the orientation of the transcription of the As resistance elements.

Similarly, operon *arsRBC* in ZAP62 with locus tags G3M80_RS09295 (103 aa), G3M80_09290 (432 aa), G3M80_09285(139 aa) showed 100 % of aminoacids identity with ArsR as a winged helix-turn-helix transcriptional regulator, ArsB and ArsC proteins of *B. altitudinis*. Whilst the products of *arsRBCDA* operon in ZAP62 with locus tags G3M80_12335 (120 aa), G3M80_12330 (432 aa), G3M80_RS12325 (139 aa), G3M80_12320 (125 aa), and G3M80_RS12315 (589 aa) showed a high percentage of identity not only with members of *B. pumilus* group such as *B. altitudinis* and *B. pumilus*, but also with species as *B. paralicheniformis*, *B. licheniformis, B. swezeyi,* and *B. atrophaeus*. Additionally, a third ArsR and ArsB were found in ZAP62 with locus G3M80_09305 (107 aa) and G3M80_RS01765 (447 aa) identified as a metalloregulator ArsR/SmtB family transcription factor and an arsenical efflux pump membrane protein.

Furthermore, to have an overview of the phylogenetic relationships of the *ars* genes with other species, protein sequences were used to infer evolutionary relationships (Figure 6). Thus, phylogenetic analysis revealed that the ArsR sequences belong to families of metalloregulators ArsR/SmtB and winged helix-turn-helix transcriptional regulators. This type of transcriptional regulators can be found in different species of the *Bacillus* genus, but also other genera, such as *Staphylococcus*.

**Figure 6.**
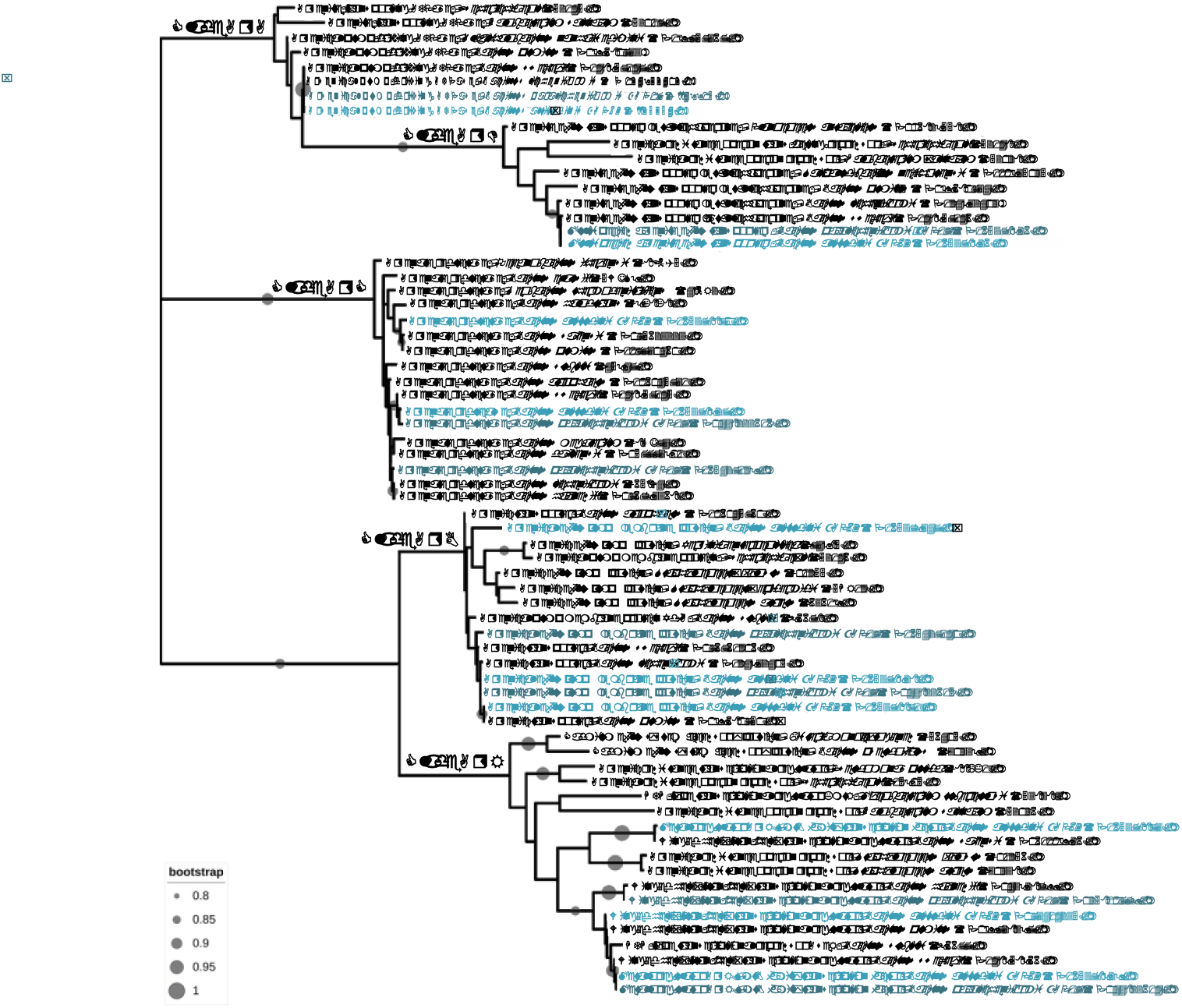
Phylogenetic relationships of ars proteins found in ZAP strains and related sequences. The phylogenetic tree was constructed using the Maximum Likelihood method and bootstrap support of 1000 replications. The inner circles show replications greater than 80%. Reference sequences are indicated in parentheses.

Arsenic resistance systems are widely distributed in prokaryotes (Fekih et al., 2018) and *ars* operons can be as simple as the *arsRB* operon in species like *Ferrolasma acidarmanus* and *Leptospirillum ferriphilum* (Gihring et al., 2003; Tuffin et al., 2006), or as complex as in *Leclercia adecarboxylata* As3-1 with four gene clusters involved phoUpstBACS, arsHRBC, arsCRDABC, ttrRSBCA (Han et al., 2019) (Figure 5).

ArsR–SmtB family members possess a highly conserved DNA recognition helix-turn-helix (HTH) motif that binds to their operator/promoter region repressing the expression of operons in absence of metal ions such As, Sb, Bi, Zn, Cd, Pb, Co, Ni, Cu, and Ag (Saha et al., 2017; Antonucci et al., 2018; Prabaharan et al., 2019). In this way that repressors found in the ZAP17 and ZAP62 genomes could be capable of showing resistance to other metal ions.

ArsB sequences showed high similarities not only with *Bacillus* species but also with other ArsB sequences in genomes of *Staphylococcus*, *Yersinia,* and *Escherichia*. Interestingly G3M80_09305 and G3M80_RS01765 genes, the third ArsR (WP 165378873.1) and ArsB (WP 165379347.1) found in ZAP62 were the sequences that showed the greatest similarities with other species different from the *Bacillus* genus. On the other hand, ArsC sequences were identified as thioredoxin type, with their three redox-active cysteine residues Cys10, Cys82, and Cys89 which are critical for the enzymatic activity of ArsC (Li et al., 2007) (Supplementary Figure 1). ArsC proteins showed high similitudes among *Bacillus* species. Besides, ArsD and ArsA proteins were identified as a metallochaerone and an ATPase that works in conjunction with ArsB enhancing the rate of arsenite extrusion (Lin et al., 2006; Garbinski et al., 2019). Both proteins were found closely related to other *Bacillus* species (Figure 6).

In addition, to act as a metachaperone, ArsD functions as a weak transcriptional regulator of the ars operon (Lin et al., 2007; Ajees et al., 2011). The predicted ArsD cluster in ZAP genomes contains Cys12-13 and Cys-18 residues and lacks Cys112-113 and Cys119-120 (Supplementary Figure 1). This presence and absence of residues confer the activity of metallochaperone to ArsD (Lin et al., 2006; Ajees et al., 2011). Thus, the *arsD* product in both strains could act only as a metallocahaperone, besides phylogenetic analysis showed a distant relationship with ArsD with repressive activity such as *E. coli* (Figure 6).

High concentrations of As have probably led to exerting evolutionary pressure on the Ars operons, increasing their complexity and efficiency with the addition of genes such as AsrD and ArsA (Mukhopadhyay and Rosen, 2002; Zhu et al., 2014; Fekih et al., 2018). In this work, two *ars* operons are evidenced in the genomes of ZAP17 and ZAP62, *arsRBC,* and *arsRBCDA*. This arrangement of genetic elements of As resistance and gene redundancy has already been reported in other Bacilli (Musingarimi et al., 2010; Bhat et al., 2011).

According to Fekih et al. (2018) it is common to find multiple *ars* operons in bacteria as a result of gene horizontal transfer and duplication. On the other hand, genetic redundancy seems to be the solution to the regulation of the various As resistance genes, since it allows the differential expression of theseunder various environmental factors (Páez-Espino et al., 2014; Zhao et al., 2015; Han et al., 2019). The *arsRBC* and *arsRBCDA* operons found in ZAP17 and ZAP62 are the putative genetic elements associated with their As tolerance, such as As level found in the Araró hot springs.

### 3.4. Expression analysis of *ars* genes

To gain further insight into the functionality of *ars* operons found in ZAP17 and ZAP62, and the possible involvement of arsenite in the expression of *ars* individual genes (*arsR, arsB* and *arsR, arsB, arsD, and arsA*), semiquantitative RT-PCR assays were carried out using total RNA from ZAP17 and ZAP62 cultures untreated (0 mM) and supplemented (12 mM) with arsenite (Figure 7). The 12 mM concentration is a subinhibitory concentration of arsenite. Genomic DNA (gDNA) was used as a positive control, as well as PCR mix as negative control (data not shown). No expression changes in the 16S rRNA genes from both strains were observed. The expected amplicons, corresponding to each *ars* genes, were detected not only in the arsenite-treated cultures but also in non-treated cultures. Although it could be argued that an induced expression of the *ars* genes is noted in the presence of the metalloid, a more detailed analysis is required, such as qPCR.

**Figure 7.**
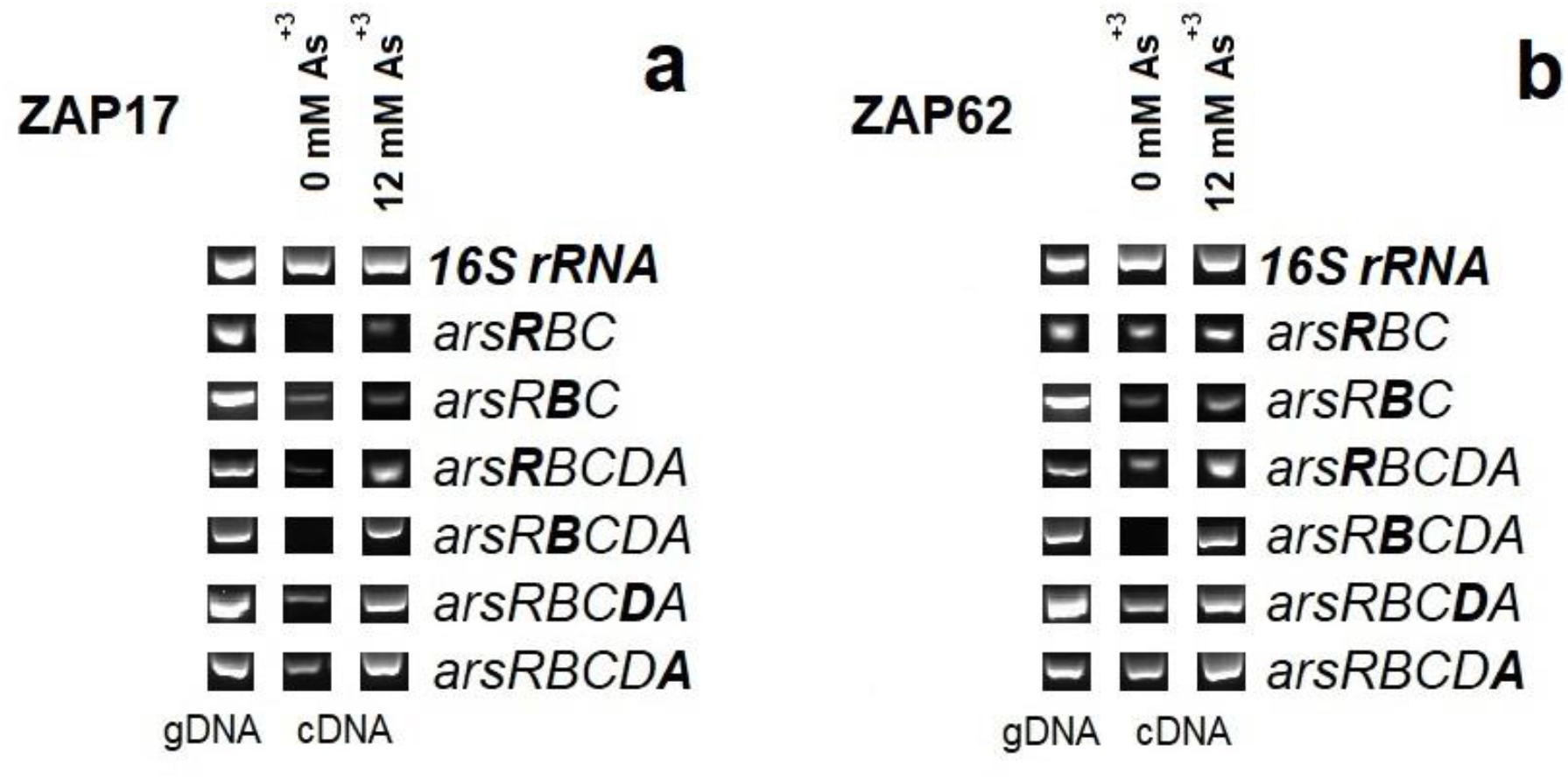
Basal and induced expression of the *ars* genes. The amplicons obtained in the gene expression assays are shown as fragments of electrophoresis gels, the bands obtained from each region were evaluated with a molecular weight marker. Section **a** shows the amplicons obtained in the ZAP17 expression assays, while section **b** shows those of ZAP62. In the figure it is also possible to appreciate the functionality of the primers using gDNA as a template. The experiments were repeated at by triplicate with similar results.

The expression of *ars* genes has also been detected in other bacterial species, such as *Burkholderia xenovorans* LB400 (Serrato-Gamiño et al., 2018), demonstrating, along with heterologous expression analysis of the ars operon, the role of such genetic elements to resist concentrations of the metalloid. However, such As-resistance is not comparable with the elevated concentrations of As highly-resistant Bacilli strains (Botes et al., 2007; Badage et al., 2020), including ZAP17 and ZAP62. In this sense, Firmicutes are regular inhabitants of hot springs with elevated concentrations of diverse heavy metals and metalloids, such as As (Das et al., 2016; Prieto-Barajas et al., 2017; Zhang et al., 2020; Oremland and Stolz, 2003). Therefore, having a functional genetic reservoir of *ars* operons in their genomes is fundamental for survival. However, other genetic elements involved in As resistance were not explored in this work, like *arsM* (which encodes an As (III) S-adenosylmethionine methyltransferase), *arsI* (a C–As bond lyase), and *arsH* (a gene encoding a methylarsenite oxidase), or other enzymes involved in ROS detoxification and tolerance (Yang and Rosen, 2016).

## 4. Conclusion and future perspectives

*Bacillus paralicheniformis* ZAP17 and *Bacillus altitudinis* ZAP62 harbor two functional *ars* operons, which allow them to resist elevated concentrations of arsenate or arsenite, a metalloid commonly found in aquatic environments, like the hot spring microbial mats from Araró, México. Thus, these Bacilli strains could be an option for future bioremediation strategies, as well as biocontrol agents and plant growth promoters, particularly, in agricultural soils contaminated with metals.

## 5. Acknowledgements

The authors thank Dr. Carlos Cervantes for supplying the arsenate reagent and Julie Hernandez-Salmeron for technical support.

## Suppl. Data

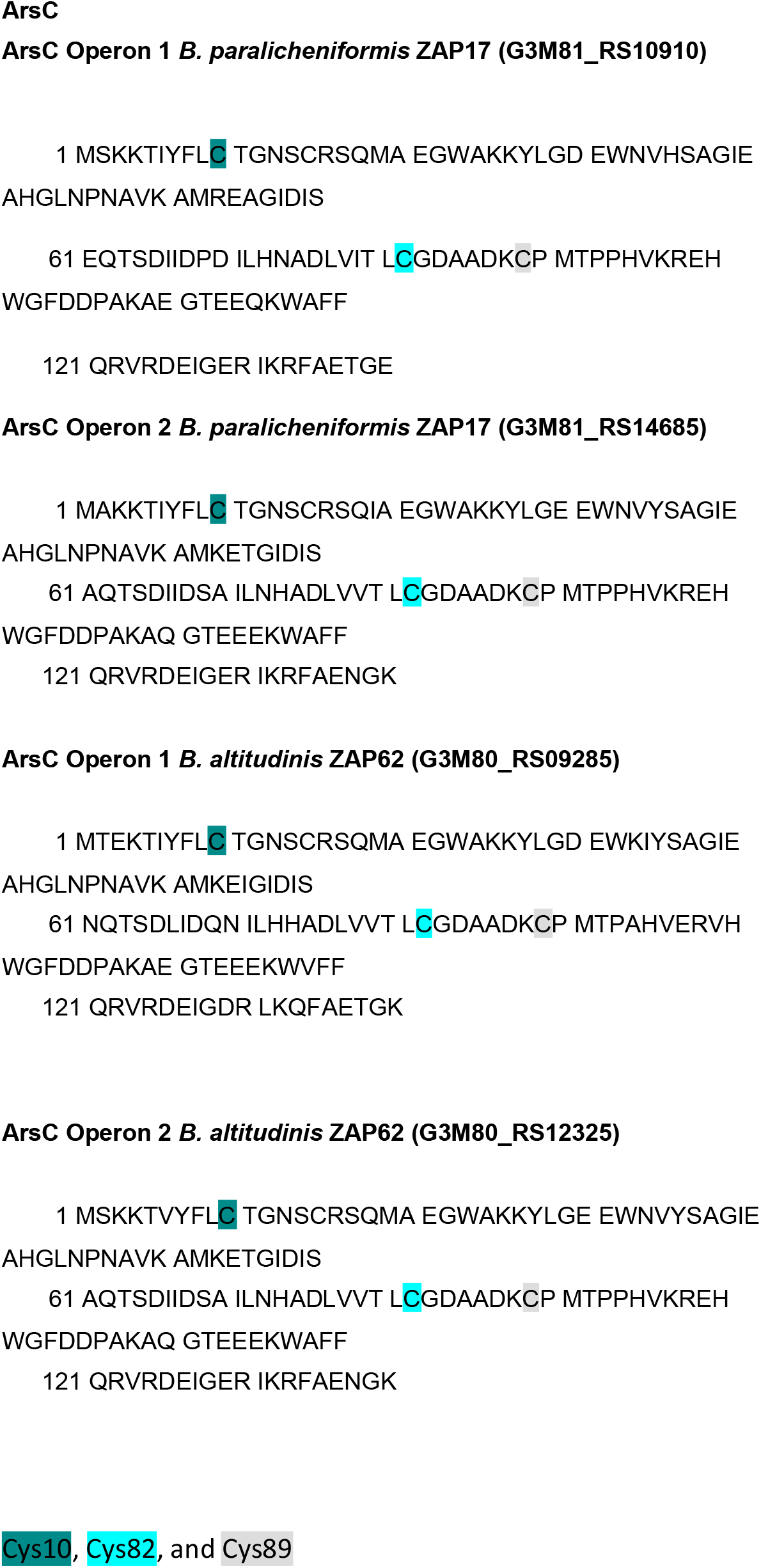

## Notes

### Competing Interest Statement

The authors have declared no competing interest.

